# Poor Generalization by Current Deep Learning Models for Predicting Binding Affinities of Kinase Inhibitors

**DOI:** 10.1101/2023.09.04.556234

**Authors:** Wern Juin Gabriel Ong, Palani Kirubakaran, John Karanicolas

## Abstract

The extreme surge of interest over the past decade surrounding the use of neural networks has inspired many groups to deploy them for predicting binding affinities of drug-like molecules to their receptors. A model that can accurately make such predictions has the potential to screen large chemical libraries and help streamline the drug discovery process. However, despite reports of models that accurately predict quantitative inhibition using protein kinase sequences and inhibitors’ SMILES strings, it is still unclear whether these models can generalize to previously unseen data. Here, we build a Convolutional Neural Network (CNN) analogous to those previously reported and evaluate the model over four datasets commonly used for inhibitor/kinase predictions. We find that the model performs comparably to those previously reported, provided that the individual data points are randomly split between the training set and the test set. However, model performance is dramatically deteriorated when all data for a given inhibitor is placed together in the same training/testing fold, implying that information leakage underlies the models’ performance. Through comparison to simple models in which the SMILES strings are tokenized, or in which test set predictions are simply copied from the closest training set data points, we demonstrate that there is essentially no generalization whatsoever in this model. In other words, the model has not learned anything about molecular interactions, and does not provide any benefit over much simpler and more transparent models. These observations strongly point to the need for richer structure-based encodings, to obtain useful prospective predictions of not-yet-synthesized candidate inhibitors.

## Introduction

Predicting drug-target interactions (DTIs) quantitatively and accurately is a key task in guiding drug discovery. The human kinome holds great promise as a source of drug targets given their role in numerous cellular processes and inherent druggability [1,2]. Kinase inhibitors already have a strong record as pharmaceuticals, with more than 70 unique chemical entities receiving United States Food and Drug Administration (FDA) approval and many more in development [3]. Kinase inhibitor libraries, however, are often large due to the diversity of chemical space containing canonical hinge-binding motifs [4,5]. Accordingly, combing through large kinase inhibitor libraries using traditional high-throughput screening techniques can be time consuming and costly [6], presenting a natural opportunity for contributions from complementary *in silico* techniques. For this reason, computational techniques that can be used to screen chemical libraries for kinase inhibitors would have great impact on biomedical research with potential applications to diseases including cancer [7], rheumatoid arthritis [8], and many more.

Machine learning algorithms have presented themselves as a potential tool for screening large chemical libraries. In the past, DTI prediction was approached as a binary classification problem that differentiated between interacting and non-interacting drug-target pairs [9], but recent advances in deep learning have allowed algorithms to treat DTI prediction as a regression task predicting binding affinity values such as K_d_, K_i_, or IC_50_ [10-12]. Deep learning is particularly well-suited for this task due to its ability to discover intricate structures in high-dimensional data [13]. These models are trained by minimizing an objective function that measures the error between its predictions and the true values by tuning its internal parameters [13]. The recent successes of Convolutional Neural Networks (CNNs) in image comprehension tasks have caught the attention of computational biologists [12] due to their ability to extract highly correlated local motifs [13]. Intuitively, this is useful in understanding chemical structures such as hydrogen bond donor/acceptor and aromatic structures as motifs that are invariant to location. Otherwise put, features such a benzene ring that is on different parts of two unique molecules can be recognized as the same substructure. Already these tools have already been widely used to assist in the computational screening of kinase inhibitor libraries [14,15].

Accordingly, numerous models have since been reported that seek to use deep neural networks for accurately predicting DTI of a given drug-target pair [16]. Drugs and targets are parametrized as inputs for the machine learning models in a variety of ways. Features are extracted from drugs using SMILES (Simplified Molecular Input Line Entry System) strings [12,17] and PubChem structure clustering similarity matrices [10,11]. Methods have also been developed to extract features from target proteins including using protein sequences directly [12], or using Smith-Waterman protein similarity matrices [10,11]. Meaningful feature extraction is of paramount importance to machine learning-based DTI prediction, as it allows for the model to understand the biophysical bases of kinase-inhibitor interactions. It is therefore surprising, in a sense, that protein sequences are sufficient to predict ligand binding, because the strengths of the intermolecular interactions depend sensitively on subtle details of the protein structure.

While many existing models report impressive performance in benchmark experiments, the ability of these models to generalize to previously unseen information remains unclear. Performance in this context is extremely relevant for real-world application of these models, since their value is expected to derive principally from prediction of inhibitors that have not yet been characterized yet – or even synthesized, ideally. To explore this question, we build a CNN-based model that takes protein sequences and SMILES strings to predict the binding affinity of a given drug-target pair, to serve as a direct analog of existing models in the scientific literature. As expected, performance of this model is also equivalent to existing models, motivating us to move forward and test this representative model’s ability to generalize to previously unseen data. Our findings demonstrate that randomly splitting data into training, validation, and test sets – which underlies most published studies [11,12,17-23] – introduces redundancy between training and test sets encouraging the model to memorize kinase phylogeny and match chemical analogues to make predictions instead of extracting meaningful features from SMILES strings and kinase protein sequences.

## Methods

### Dataset Preparation

We evaluated our model on four different datasets that have been previously used for DTI prediction studies [10-12,18,19,24-26]. We refer to these datasets by the last names of their respective first authors: the Anastassiadis dataset [27], the Christmann-Franck dataset [28], the Davis dataset [29], and the Elkins dataset [30].

The Anastassiadis dataset screens 178 commercially available kinase inhibitors against 300 protein kinases [27], after excluding kinase mutants. Kinase information is converted from protein sequence representations to 85-character long KLIFS sequences, corresponding to the 85 residues in kinases’ structurally-conserved active site at which ATP-competitive inhibitors bind [31,32]. Inhibition in this dataset (for each inhibitor/kinase pairing) is reported as percent activity of the inhibited kinase relative to the same kinase’s activity in the absence of inhibitor. We therefore standardize the bioactivity labels by converting percent inhibition to pIC_50_ values to align the range of training labels with that of the other datasets we use. We perform this conversion first by adjusting percent inhibition values such that those above 98% are recorded as 98% and those below 2% are recorded as 2%. Assuming a Hill slope of 1 (competitive inhibition at a single site), we then estimate half maximal inhibitory concentration through the following equation:

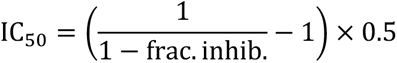

where the fraction of inhibition is the decimal value of percent inhibition. Molar IC_50_ is then further converted into pIC_50_ by the following equation:

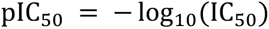

These pIC_50_ values are used as bioactivity labels for training and testing the model. We further filter the dataset to exclude datapoints where the SMILES string is more than 90 characters long to eliminate three excessively large outlier compounds. This results in 35,350 datapoints in the processed dataset.

The Christmann-Franck dataset complies bioactivity data from a number of datasets previously published in the scientific literature reporting data for 2,106 inhibitors and 482 kinases containing 356,908 datapoints [28]. This dataset is extremely large and comprehensive, incorporating data reported in the widely used KIBA dataset among others. As this dataset compiles data from a variety of sources that report bioactivity differently, Christmann-Franck and coworkers standardize bioactivity values to pAct. We filter this dataset by removing kinase mutants and non-human kinases, remove the 15 inhibitors with SMILES strings of more than 90 characters, and convert kinase data to KLIFS sequences. After processing, the dataset contains 210,679 datapoints.

The Davis dataset screens 72 kinase inhibitors against 442 kinases covering more than 80% of the human kinome [29]. In line with our other data processing techniques, we transform given dissociation constant K_d_ to pK_d_ through the following formula:

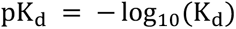

Finally, we convert kinases to KLIFS sequences and remove datapoints for 3 inhibitors where the SMILES string is more than 90 characters long. The processed dataset contains 22,839 datapoints.

The Elkins dataset is a thorough characterization of the Published Kinase Inhibitor Set (PKIS) that selectively screens 367 inhibitors against 224 recombinant kinases [30]. First, the dataset is filtered to remove any interactions with an (unphysical) dissociation constant less than or equal to 0. After converting kinase information to KLIFS sequences, removing datapoints with SMILES strings of more than 90 characters, and converting K_d_ to pK_d_ values, 48,468 datapoints remain. **Table 1** summarizes the final processed datasets used in our experiments.

**Table 1:** Summary of datasets used in our study. The bioactivity label, number of unique kinase and inhibitors, and kinase-inhibitor interaction pairs after processing.

| Dataset name | Bioactivity label type | Number of kinases represented | Number of distinct inhibitors | Total number of inhibitor/kinase interactions |
| --- | --- | --- | --- | --- |
| Anastassiadis | pIC<sub>50</sub> | 202 | 175 | 35,350 |
| Christmann-Franck | pAct | 281 | 2,091 | 210,679 |
| Davis | pK<sub>d</sub> | 329 | 69 | 22,839 |
| Elkins | pK<sub>d</sub> | 185 | 327 | 48,468 |

### Selection of Training, Validation, and Test Sets

To train and evaluate our model’s capacity to generalize to previously unseen data, we use two different methods of splitting our dataset into training, validation, and test sets. We refer to these methods as “Standard Split” and “Split by Inhibitor”. *Standard Split* uses the typical best-practice of randomly spitting all available datapoints 80%/10%/10% into the training, validation, and test sets, respectively. This represents the standard approach in machine learning, encompassing nearly all past studies that attempt to use machine learning methods to predict kinase inhibitor binding affinities [11,12,17,18]. *Split by Inhibitor* involves selecting 10% of the ligands and moving all datapoints corresponding for these compounds into the test set, and selecting a separate 10% of the ligands for the validation set, and leaving the remaining 80% as the training set. Thus, all points in the validation or test set correspond to an inhibitor that the model has never before seen in the training set: this in turn allows us to evaluation how well a model generalizes to inhibitors previously unseen in the training set. Both strategies are summarized in **Figure 1**.

**Figure 1:**
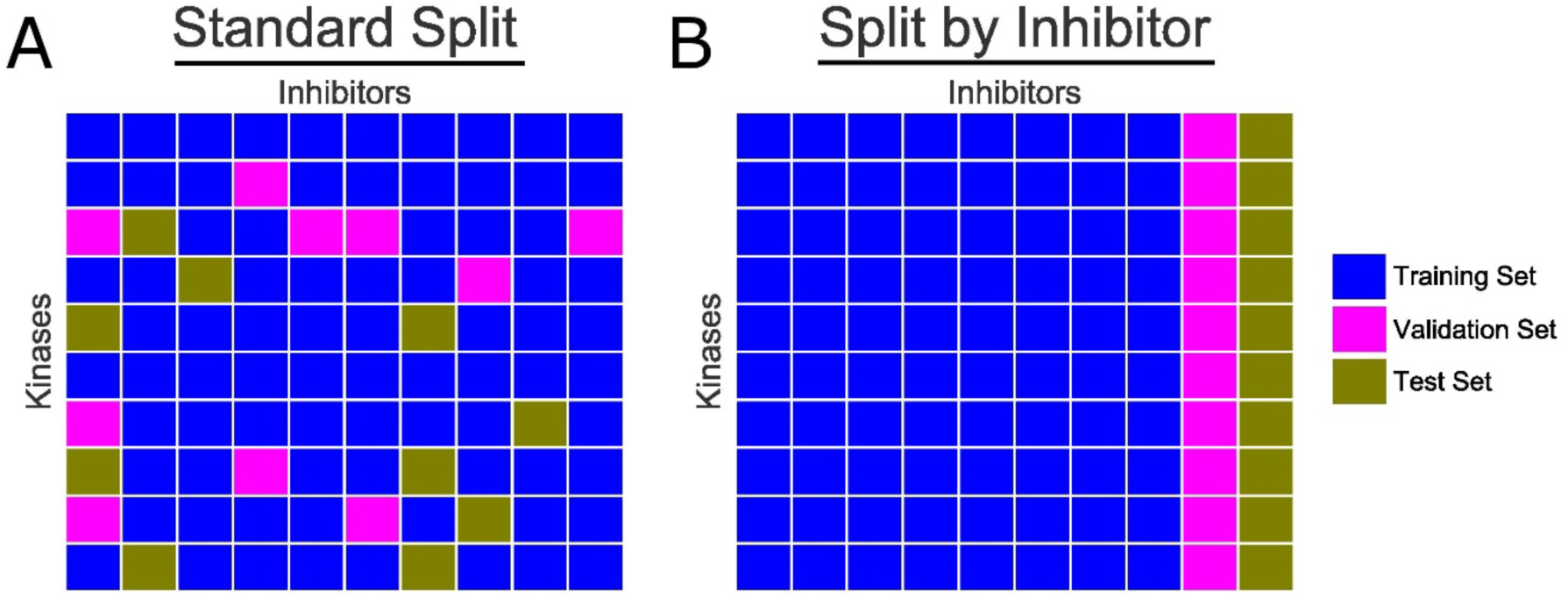
Schematic representation of dataset splitting approaches. Kinases are listed along the vertical axis, and inhibitors along the horizontal axis. Each cell represents a unique kinase-inhibitor interaction. **(A)** “Standard Split” randomly puts each data points into either the training set (*blue*), validation set (*magenta*), or test set (*gold*). **(B)** “Split by Inhibitor” holds back from the training set all data points from several of the inhibitors, such that the test set (and validation set) comprises inhibitors that have not been seen by the model during the earlier (training) steps.

### Machine Learning Model Architecture

In our study, we treat DTI as a regression problem by having the model predict the binding affinity (label) of a given kinase-inhibitor pair. In our model, we use the power of CNNs that have seen great successes in a wide variety of pattern recognition tasks [13]. CNNs are structured as a series of convolutional layers and pooling layers. These are especially suited to capture local features with the help of filters and merge semantically similar features through pooling layers [13]. CNNs have already gained popularity in the cheminformatics world as they hierarchically decompose inputs so that the network recognizes higher level features while maintaining spatial relationships [33].

Our model takes 1D representations of both kinases and ligands – through KLIFS sequences and SMILES strings – as inputs. KLIFS sequences and SMILES strings are then encoded into integer vectors of length 85 and 90 respectively, with a provision for missing amino acids in the KLIFS sequences. These integer vectors are then passed through an embedding layer that transforms the KLIFS and SMILES vectors into dense vectors of fixed size. These embedded vectors are then fed into separate CNN blocks. In each block, three consecutive 1D convolutional layers are used with an increasing number of filters followed by a pooling layer. The outputs of these CNN blocks are then concatenated and fed into a dense neural network that predicts the DTI of a given kinase-inhibitor pair. We use 1024 nodes in the first two dense layers, 512 layers for the third, followed by the output node. To prevent model overfitting, we use dropout layers of rate 0.1 in both dense and convolutional layers [34]. The model was trained over 300 epochs with a batch size of 256. Following our tuning practices, we use Rectified Linear Unit (ReLU) as our model’s activation function [13] and Adam as our optimizer [35] with a learning rate of 0.001 to minimize the mean squared error (MSE) error function.

Our model has adjustable hyperparameters such as the number of CNN filters, filter length in each CNN block, number of dense layers, nodes in each dense layer, dropout rate, and learning rate. Starting from parameter values from DeepDTA [12], we performed a systematic tuning of these parameters by searching for best performance on test data. These experiments demonstrated that the parameters used in DeepDTA provided the most consistent results. Our replication of this framework was enabled by the authors of this study making their code readily available.

Our model is implemented in Keras [36] with a TensorFlow backend [37]. Our model architecture is illustrated in **Figure 2**.

**Figure 2:**
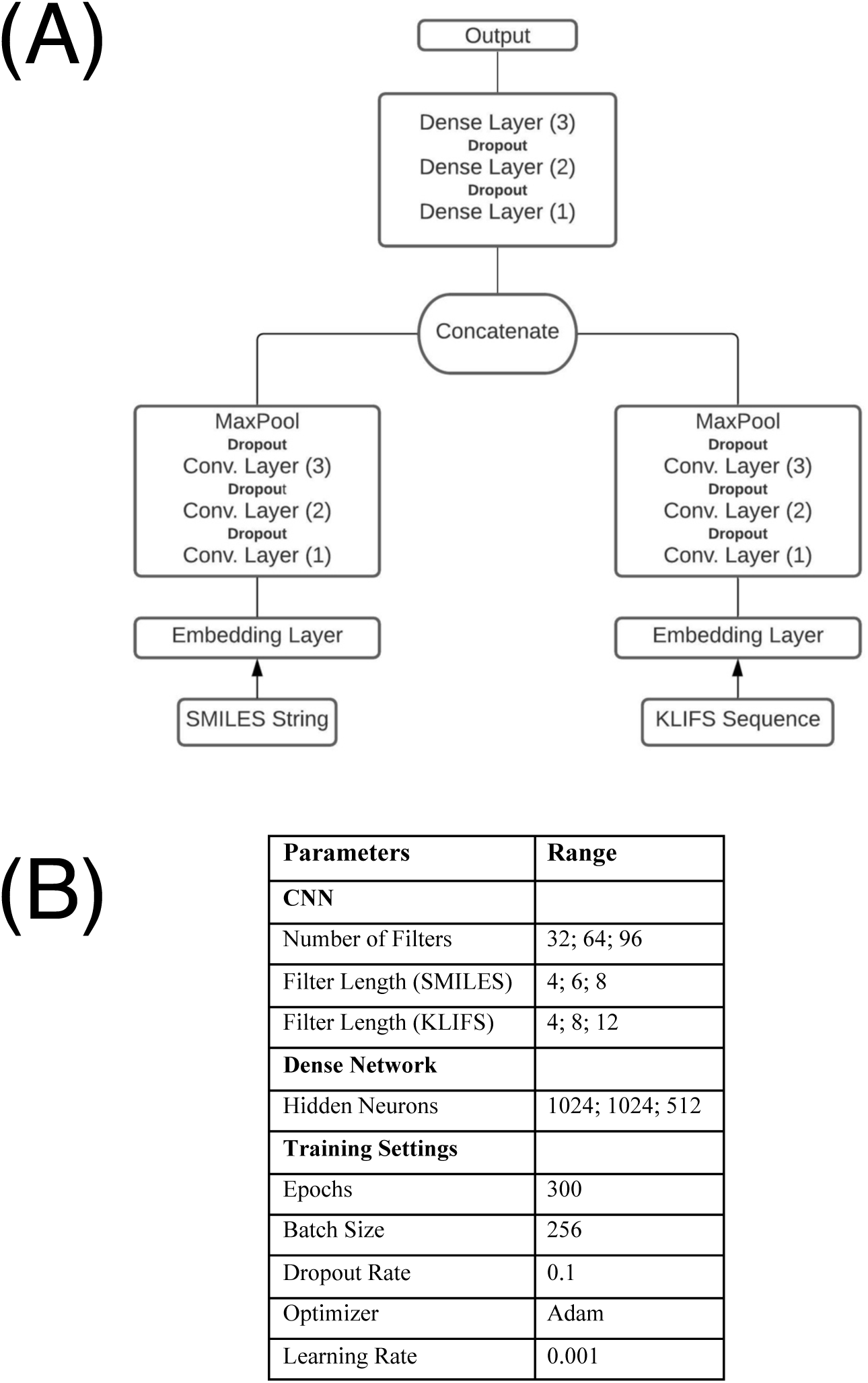
Schematic of the CNN model used in this study. **(A)** The model takes 1D SMILES and KLIFS integer vectors and transforms them into dense vectors of fixed size. These dense vectors are passed into CNN blocks, each with 3 convolutional layers with one max pooling layer at the end. The outputs of the CNN blocks are concatenated and fed into a dense network of three layers to predicts the binding affinity (pActivity) of a given kinase-inhibitor pair. **(B)** Experimental tuning led to selection of the parameters shown, which are similar to those used in others’ related models.

### Junk SMILES Experiment

To probe for information leakage caused by randomly split cross-validation data, we also carried out an experiment in which we replaced the SMILES string information for each inhibitor with a random string of equivalent length to the inhibitor’s original SMILES string. All data for a given inhibitor is thus associated with a meaningless (but constant) placeholder SMILES string that encodes no information about the inhibitor. These randomized strings are generated by arbitrarily replacing each character in each inhibitor’s SMILES string with another character in the SMILES dictionary. These strings do not encode any meaningful chemical information, but rather they simply serve as “tokens” that assign a distinct identity to each inhibitor. The model is then trained with KLIFS sequences and these character strings in place of SMILES strings. Data is split between the training, validation, and test sets using the Standard Split approach described above.

### 2D Molecular Comparisons

We also designed an additional experiment to further evaluate potential information leakage arising from our model making predictions based on memorization of closely related chemical analogues in the training set. We use various tools to calculate 2D chemical similarity such as the Molecular Access System (MACCS) in RDKit [38] and other methods described in a comparative study of SMILES-based similarity kernels [39]. The MACCS system represents the presence/absence of 166 predetermined chemical features and calculates the similarity between two compounds using the Dice coefficient [40]. The maximum common substructure (MCS) method examines the common atoms and bonds between two molecules to determine a similarity score. The edit distance method uses the number of edits needed to convert one string into another as a measure of similarity. The Combination of LCS models (CLCS) similarity measure based on a combination of normalized longest common subsequence and maximal consecutive longest common subsequence [41]. The LINGO chemical similarity measure examines the q-character substrings of SMILES text and have been previously used as quantitative structure-property relationship models [41]. The substring method defines similarity as the inner product of the frequencies of all substrings of two or more characters [39]. The TF similarity measure compares the frequency of four-character substrings between two compounds [39]. Finally, the TF-IDF frequency combines the TF similarity measure by multiplying it with the inverse document frequency [39]. Using this information, we take the binding affinity values of this closest training set compound as determined by each of the similarity measures and that compound’s binding affinity values as the predictions of a “model”.

### Statistical Analysis

For statistical analyses, we used Lifelines [42] to calculate the concordance index and the SciPy scientific computing package for other analyses [43]. These libraries are implemented in Python.

## Results

### CNN Model

We evaluate the performance of our model on the four datasets (Anastassiadis, Christmann-Franck, Davis, and Elkins) using each of the train/validation/test splitting approaches described above. This model incorporates both CNN and deep learning architectures to predict the binding affinity of a given kinase-inhibitor pair using SMILES strings and a kinase’s KLIFS protein sequence. Of most significant note to us are studies that leverage both SMILES strings and protein sequences to predict DTI [12,17,18]. However, it is also important to compare the success of our approach against other models reported in the literature – including those that use different parametrizations of kinases and inhibitors. Previous models have elected to use concordance index as a metric to evaluate model predictions which evaluates the extent to which predicted binding affinities of kinase-inhibitor pairs are predicted in the same order as their true values [44]. This metric is especially suited to virtual screening as it measures the ranks of a model’s predictions against the true ranks of test set data. We summarize the performance of several such models in **Table 2**.

**Table 2:** Performance of representative models from the recent literature, and of the analogous model that we use in this study. Both KronRLS and SimBoost use Smith-Waterman and PubChem similarity matrices to featurize proteins and inhibitors, whereas DeepDTA uses a CNN. Performance is reported as the concordance index (CI) and the mean square error (MSE); a model with good performance should have high concordance index (CI) and low mean square error (MSE).

| Model | Dataset | Protein Representation | Ligand Representation | CI | MSE |
| --- | --- | --- | --- | --- | --- |
| KronRLS | KIBA | S-W → Dense Network | PubChem → Dense Network | 0.782 | 0.441 |
| SimBoost | KIBA | S-W → Dense Network | PubChem → Dense Network | 0.836 | 0.222 |
| DeepDTA | KIBA | CNN | CNN | 0.863 | 0.194 |
| KronRLS | Davis | S-W → Dense Network | PubChem → Dense Network | 0.782 | 0.379 |
| SimBoost | Davis | S-W → Dense Network | PubChem → Dense Network | 0.872 | 0.282 |
| DeepDTA | Davis | CNN | CNN | 0.878 | 0.261 |
| *Our model* | *Davis* | *CNN* | *CNN* | *0.896* | *0.177* |

We tested our models using the model checkpointed with the lowest validation loss for each dataset technique on each of the four datasets we included in our study (**Figure S1**). To facilitate comparison with previously published models, we report the concordance index (CI) and mean square error (MSE) in addition to the Pearson’s correlation coefficient (R). Initial experimental results for our model are presented in **Figure 3**, and these are summarized in **Table 3**.

**Figure 3:**
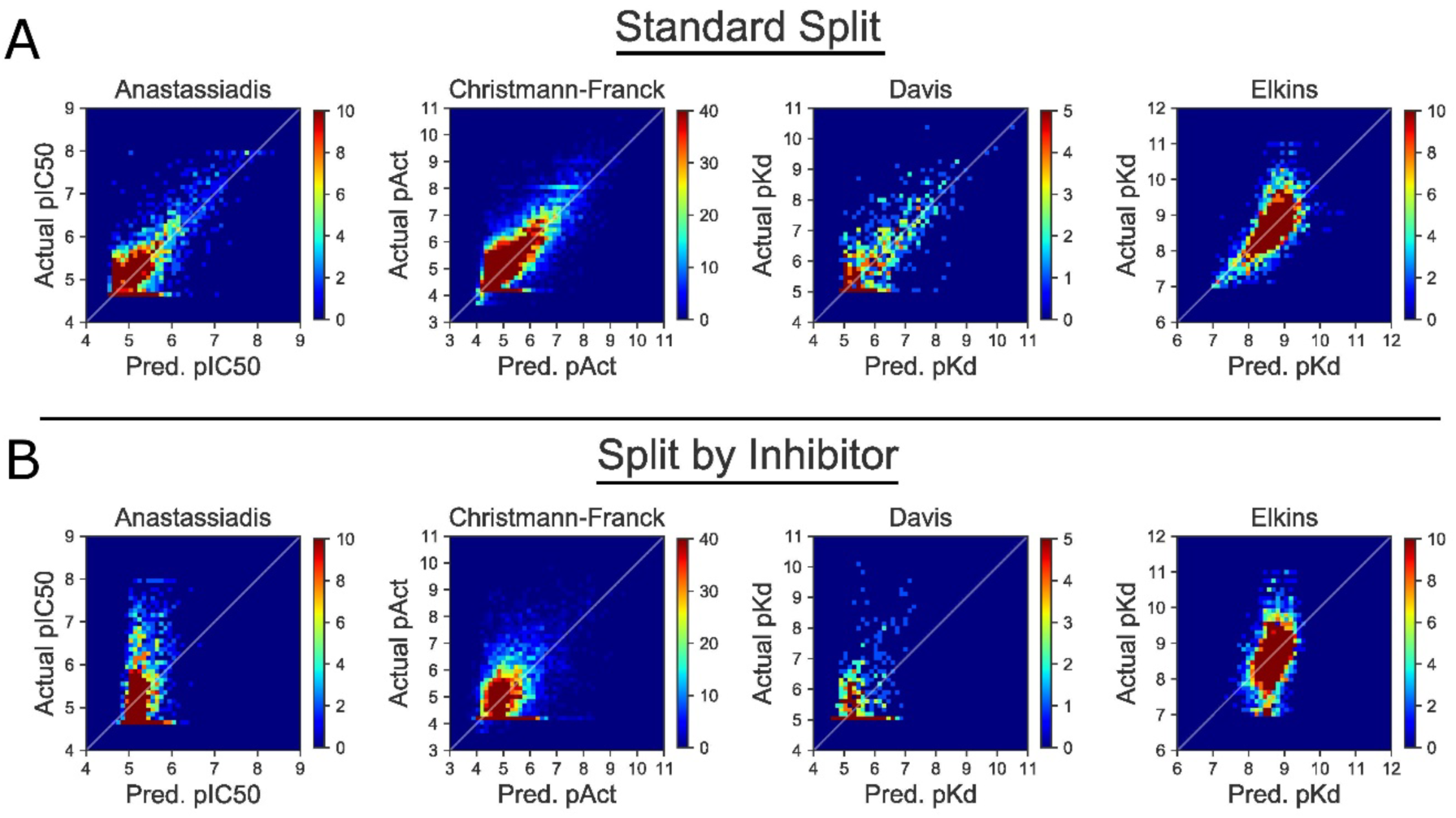
Using “Split by Inhibitor” diminishes model performance relative to “Standard Split”. For each point in the test set, we plot the actual pActivity values (ground truth) and predicted values (from the CNN). Density plots are presented for clarity, and the corresponding scatterplots are included as **Figure S2**. Clear correlations between actual and predicted values are evident with “Standard Split” (*top*), but these are lost when the model is trained using “Split by Inhibitor” (*bottom*). Quantitative performance measures from this experiment are included as **Table 3**.

**Table 3:** Using “Split by Inhibitor” diminishes model performance relative to “Standard Split”. When “Standard Split” is used, our representative model yields equivalent performance to previously reported methods that also use this splitting approach. When using “Split by Inhibitor”, the model performance is dramatically deteriorated in all four testcases.

| Dataset Splitting Approach | Dataset | CI | MSE | Pearson R |
| --- | --- | --- | --- | --- |
| Standard Split | Anastassiadis | 0.763 | 0.173 | 0.785 |
|  | Christmann-Franck | 0.799 | 0.297 | 0.828 |
|  | Davis | 0.896 | 0.177 | 0.859 |
|  | Elkins | 0.765 | 0.147 | 0.685 |
| Split by Inhibitor | Anastassiadis | 0.631 | 0.390 | 0.377 |
|  | Christmann-Franck | 0.634 | 0.669 | 0.459 |
|  | Davis | 0.694 | 0.226 | 0.409 |
|  | Elkins | 0.650 | 0.258 | 0.396 |

First, we find that our Standard Split model (**Figure 3a**) performs at a similar level to those previously described in the scientific literature with a comparable CI of approximately 0.8 and exhibiting relatively low mean squared error (MSE). It should be noted that MSE varies with the size and values of data in a dataset and hence should not be used to compare performance of models across different datasets. Overall, this demonstrates the near equivalence of these examples of analogous models (**Table 3**).

However, we also observe that performance deteriorates significantly when trained using data that is organized using “Split by Inhibitor” (**Figure 3b**). For all four testcases the CI and Pearson’s correlation coefficient are lower than those for the Standard Split model, and the MSE is higher. Together these indicate an inability to generalize to novel data, because the model is unable to make accurate predictions when presented with previously unseen inhibitors.

### Junk SMILES

To test for the presence of information leakage from kinase data, we train our model on random strings of characters in place of SMILES strings (dubbed “Junk SMILES” in the Methods section). By replacing SMILES strings with meaningless, but unique, strings of characters (i.e., by “tokenizing” the inhibitor identities), we allow our model to differentiate between inhibitors without understanding their structure. This provides an avenue for our model’s base predictions of a given kinase inhibitor pair, by using the training labels of an inhibitor’s binding affinity against closely related training set kinases to make a prediction on the kinase inhibitor pair. Our Junk SMILES model performs surprisingly well with CI and Pearson’s correlation coefficient almost equivalent to that on our Standard Split model and similar MSE. On selected metrics, our Junk SMILES model even outperforms the Standard Split model. On the Anastassiadis dataset, the Junk SMILES has a CI 0f 0.768 over the CI of 0.763 in Standard Split. The almost-equivalent performance of the Junk SMILES model indicates the presence of information leakage that solely arises from the model learning kinase phylogenetic relationships, and of memorizing specific tokens (inhibitors’ SMILES strings) rather than learning to parse their meaning. Experimental results and scatterplots of our predictions are presented in **Figure 4** and **Table 4**, respectively.

**Figure 4:**
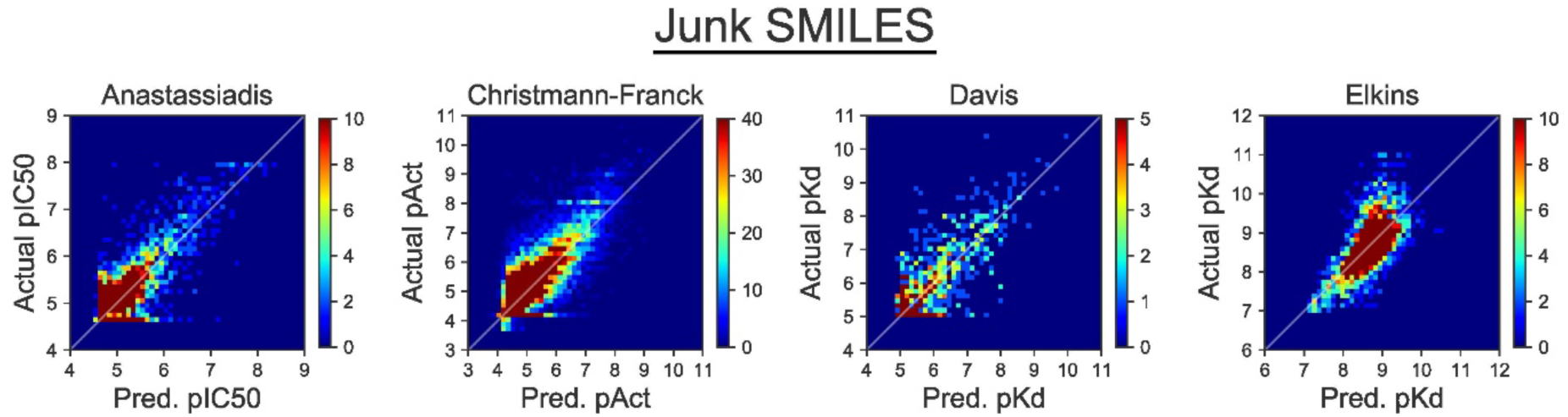
Using a tokenized inhibitor encoding does not diminish model performance. In this experiment, the identity of each inhibitor is represented by a “Junk SMILES” string that serves as a meaningless identifier. The apparently impressive performance of this model shows that this performance can be achieved by a model that does not actually parse the chemical meaning of the inhibitors. Density plots are presented for clarity, and the corresponding scatterplots are included as **Figure S3**. Quantitative performance measures from this experiment are included as **Table 4**.

**Table 4:**
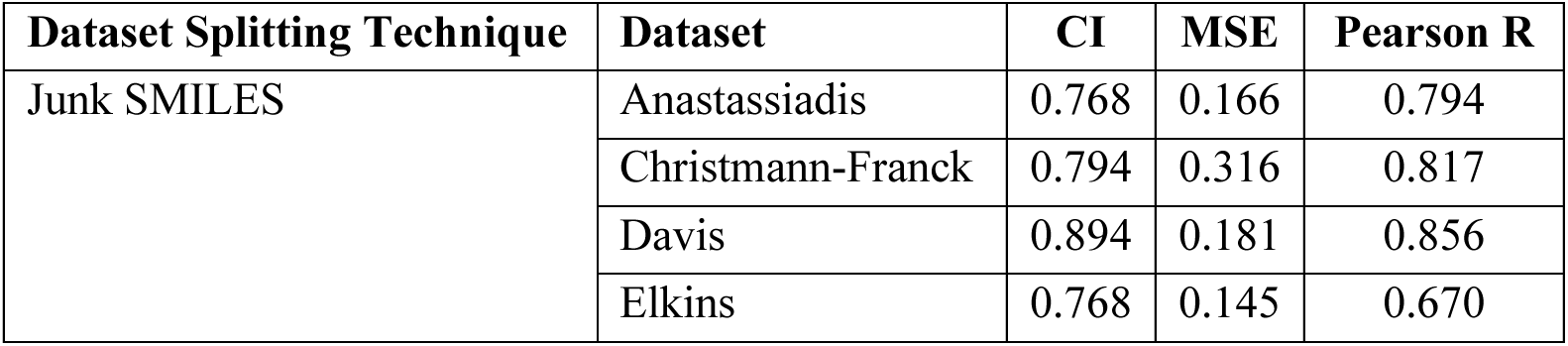
Using a tokenized inhibitor encoding does not diminish model performance. Despite lacking the information that encodes the chemical structure, Junk SMILES strings do not lead to worse model performance when “Standard Split” is used, confirming the extent to which the apparent performance of published models may be reliant on informational leakage.

### 2D Molecular Comparisons

If our model’s predictive capacity arose solely from the underlying kinase phylogeny, one would expect performance when using the Split by Inhibitor scheme to be no better than random. **Table 3** shows that this was not the case, however, implying that some other additional form of information leakage may also be present.

When carrying out splitting using the “Split by Inhibitor” approach, the particular inhibitors that are selected for inclusion in the test set are chosen randomly. Because of the nature of the experiments that underlie the available data, however, there are often close chemical analogs of these compounds in the training set (or the validation set). Given that shared chemical scaffolds typically contain a recognizably similar subsection of their SMILES strings, we hypothesized that our CNN model could be providing apparently useful performance for unseen inhibitors by simply repeating back the activity data for the closest chemical analog in the training set. This might yield good predictions, while still allowing the model to remain blissfully ignorant of anything relating to chemistry and molecular interactions.

To test this hypothesis, we implemented a naïve prediction model that we dubbed “Label Transfer”. When asked to provide a prediction for a particular inhibitor/kinase pairing, the model simply looks up the pActivity value for the most similar inhibitor in the training set with the kinase of interest. Thus, the model is not even required to carry out any form of interpolation: it simply repeats back the answer for the closest problem in the training set. This model essentially implements the simplistic one-nearest-neighbor (1NN) strategy that others have used to establish a comparative baseline when evaluating more sophisticated prediction methods [45,46]. For the step that identifies the most “similar” analog in the training set, we separately tested each of ten different methods for quantifying similarity of chemical structures.

Results for this experiment are presented in **Table 4**. The simple family of “Label Transfer” models that do nothing more than report back data from closest chemical analogue performs nearly identically to the Split by Inhibitor CNN model. These similar levels of performance achieved by this naïve approach strongly imply that the CNN model – when used in the Split by Inhibitor framework – is doing nothing more than remembering the most closely related compounds that were presented in the training set.

**Table 4:**
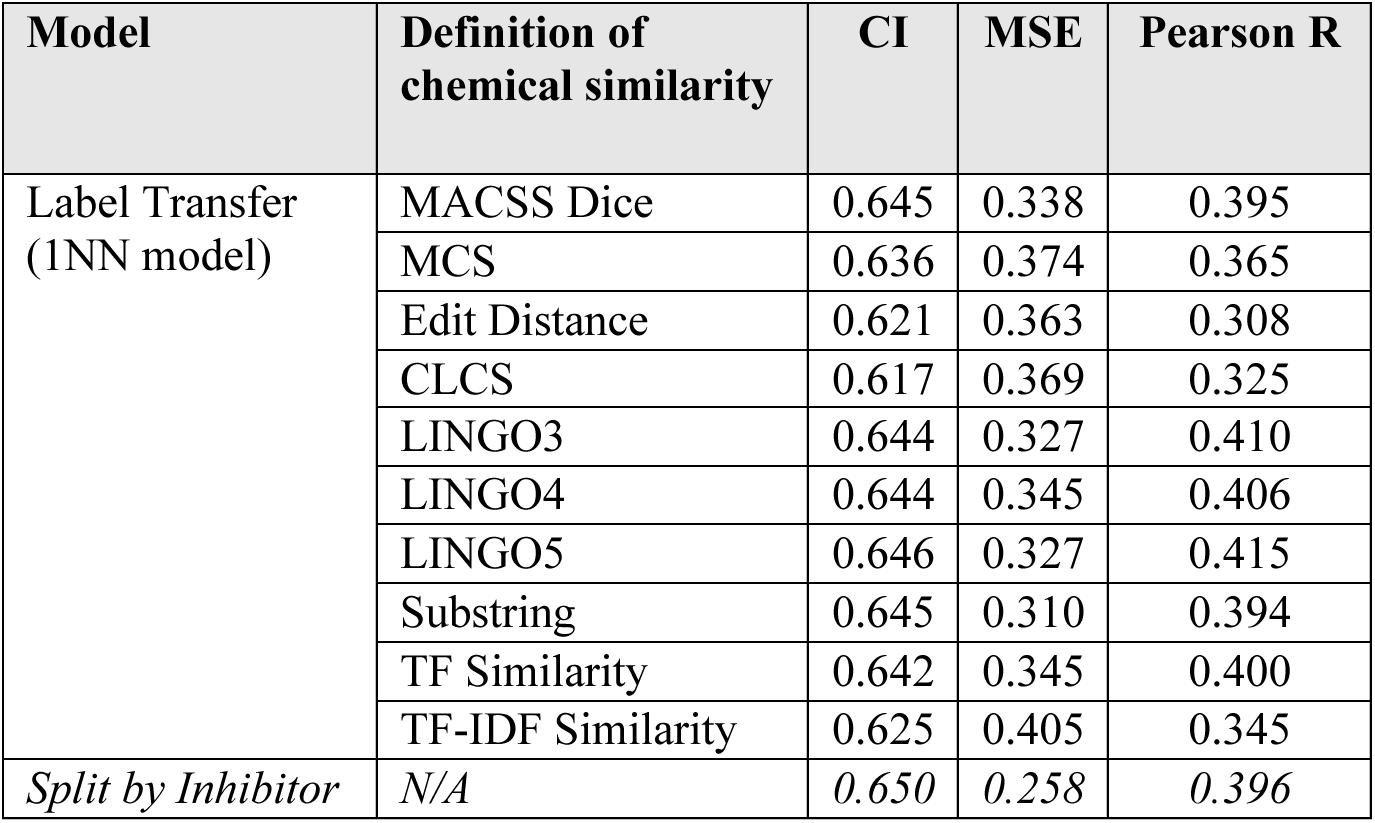
Models that simply copies activity data from the most similar chemical structure in the training set yield equivalent performance to the “Split by Inhibitor” CNN. This experiment suggests that the CNN does not necessarily generalize in the intended manner, and its performance cannot yield useful insights beyond the close analogs already included in the training set. Data in this experiment are presented for the Elkins testcase.

| Model | Definition of chemical similarity | CI | MSE | Pearson R |
| --- | --- | --- | --- | --- |
| Label Transfer (1NN model) | MACSS Dice | 0.645 | 0.338 | 0.395 |
|  | MCS | 0.636 | 0.374 | 0.365 |
|  | Edit Distance | 0.621 | 0.363 | 0.308 |
|  | CLCS | 0.617 | 0.369 | 0.325 |
|  | LINGO3 | 0.644 | 0.327 | 0.410 |
|  | LINGO4 | 0.644 | 0.345 | 0.406 |
|  | LINGO5 | 0.646 | 0.327 | 0.415 |
|  | Substring | 0.645 | 0.310 | 0.394 |
|  | TF Similarity | 0.642 | 0.345 | 0.400 |
|  | TF-IDF Similarity | 0.625 | 0.405 | 0.345 |
| *Split by Inhibitor* | *N/A* | *0.650* | *0.258* | *0.396* |

In light of these observations, it is difficult to imagine how such a model could extrapolate to make useful predictions for prospective data, such as predicting binding affinities for new compounds that have not yet been synthesized.

## Discussion

When using the “Standard Split” scheme that is typical for many machine learning applications, our benchmark CNN model yielded equivalent apparent performance to other similar recently described models. Upon closer examination, however, it becomes evident that this model is not useful at all: its performance derives solely from reporting back specific information that was present in the training set. The only true information that the model gleaned from the training data was the phylogenetic relationship between kinases, and the ability to recognize substrings in the inhibitors’ SMILES strings; though useful, both of these insights are exceedingly simple and evident from the text-based encodings presented to the model. Importantly, the model failed to *generalize* the information in the training set, which is the overarching objective for any machine learning application: successfully learning to generalize from the training set data allows a model to usefully interpolate – or even extrapolate – for the new inputs that are provided when the model is used in a realistic setting.

Even before the popularization of deep learning models in drug discovery, concern was raised about simpler machine learning models (e.g., random forests) being used as “black boxes”; in one example, it was demonstrated that this could lead to non-sensical docking scoring functions that gave back results insensitive to the docked poses it was meant to evaluate [47]. The failure of certain models to properly generalize has also been studied in the context of virtual screening, leading one group to propose that developing suitably-generalized models is hindered primarily by that fact that available experimental data sets only cover small and biased subsets of chemical space [48,49].

With respect to predicting the binding affinities of drug-like small molecules with their targets (drug-target interactions, DTIs) via deep learning, others have also recently explored several different protein families in considering the effect of various representations (encodings) of both the ligand and the protein, as well as different strategies for splitting. In agreement with the results presented at the outset of our study, these authors also found that randomly splitting datasets into training and testing folds led to near-complete data memorization and produced highly over-optimistic results [50,51].

Despite its suitability for other types of applications (where points in each data set are truly uncorrelated), there has begun a gradual recognition that random splitting is inappropriate for DTI-type prediction tasks. This has spurred some researchers to group all the data for a given inhibitor into either training or test set (which we dubbed “Split by Inhibitor”), or to group all the data for a given kinase into either the training or the test set (“Split by Kinase”) [52-55]. As we have shown here, however, this practice is insufficient to guard against information leakage, if close chemical analogs remain distributed between the training and test sets.

Rather, the experiments in our study strongly underscore the need to be very deliberate when ensuring that chemical analogs are all placed *together* in the same training/test fold, to truly avoid information leakage. The downside of this approach is that it prevents the models from learning and prospectively employing subtle details that distinguish the analogs, and it provides a very pessimistic outlook of the model performance. Instead of simply grouping all analogs together into a given training/test fold, one can instead cluster all molecules based on their similarity, then build a test set comprised of the most “distinct” member of each cluster (chemical scaffold). This strategy – described by as “neighbor splitting” or “scaffold splitting” [49,56-60] – does not entirely eliminate the information leakage that underlies “Standard Split” and “Split by Inhibitor”, but rather minimizes the value of the leaked information to best mimic a real-world scenario in which prospective inputs are not wholly independent from the training examples.

An intriguing alternative approach involves building training/test folds based on time. Provided the date at which experiments were conducted (or the date at which data became available), one can define a split such that all data acquired before a certain time are included in the training set, and data collected later represent the test set [56,57]. This “time-split” strategy is incredibly satisfying from an intuitive standpoint, because it mimics a scenario in which a model is trained based on available data, and then later applied in a prospective context to new data that did not even exist at the time of training. Gratifyingly, application of a time-split training strategy has also detected information leakage in other model frameworks, by showing that the models did not generalize to targets not available at the time of training [61,62]. It has been argued that time-splits give a more realistic estimate than the overly pessimistic estimates from “scaffold splitting”, since real-world applications will sometimes be asked to make prospective predictions for inhibitors not very dissimilar from those in the training set [16,63,64]. Moreover, there may be subtle and systematic biases if newer data come from experiments that were explicitly designed with knowledge of past results.

## Conclusions

Randomly splitting datapoints into training, validation, and test folds has long constituted a traditional best-practice in data science. However, adopting this approach for chemical structures is extremely susceptible to information leakage, because of the inherent shared activity among closely related chemical structures / closely related receptors. Accordingly, it is unequivocally necessary to adopt more thoughtful methods for splitting datasets when considering prediction problems involving drug-target interactions.

Ultimately, the splitting method should reflect the goals of the intended application for the model, and it should be carefully designed to align with the intended domain of applicability. The most rigorously extreme strategy is to exclude from the training set any points that bear any relationship whatsoever to points in test set: however, makes it extremely challenging to train the model, because some of the most potentially valuable data has been withheld from training. This in turn may lead to worse performance in real-world scenarios, where inputs may often resemble examples that would have been available in training.

Taking a broader vantage point, we suggest that the real value of any deep learning model should be evaluated relative to thoughtful control experiments. Quantitative performance measures (such as CI and MSE) are inherently sensitive to the dataset and to the similarity between training and test sets. Accordingly, these measures hold the greatest value not as a standalone number, but rather as a means to compare between models in a carefully controlled setting. By defining alternate models, one can therefore test specific hypotheses: for example, here we compared performance of a typical CNN framework against the tokenized “Junk SMILES” model and the simple 1NN “Label Transfer” model. By showing that the CNN did not out-perform either of these simpler and more transparent models, it becomes clear that the extra complexity (and opacity) of the deep learning model is not bringing additional value.

Moving forward, we are optimistic that deep learning methods will ultimately provide valuable insights for predicting binding affinities of new compounds that have not yet been synthesized, and that such models will help guide drug discovery efforts. At this point, however, it does not appear that current text-based encodings of the ligand and receptor (SMILES and protein sequence), with currently available datasets, contain sufficient information for anything more than providing trivial insights that would be evident from much simpler analysis.

## Data and Code Availability

Source code and processed datasets are freely available on the karanicolaslab GitHub repository at: https://github.com/karanicolaslab/kinbadl_klifs_SMILES_project.

## Supplemental Information

The training and validation loss plots for all datasets and approaches are shown in **Figure S1**. **Figure S2** and **S3** present the data from **Figure 3** and **Figure 4** as scatterplots rather than heatmaps.

## Acknowledgements

We thank Dr. Albert J. Kooistra for making KLIFS sequences for 497 kinases available to us. We thank Austin Clyde for assistance and useful discussions.

This work was supported by grants from the National Science Foundation (CHE-1836950) and the NIH National Institute of General Medical Sciences (R01GM141513). This research was also funded in part through the NIH/NCI Cancer Center Support Grant P30 CA006927.

This work used the Extreme Science and Engineering Discovery Environment (XSEDE) allocation MCB130049, which is supported by National Science Foundation grant number 1548562. This work also used computational resources through allocation MCB130049 from the Advanced Cyberinfrastructure Coordination Ecosystem: Services & Support (ACCESS) program, which is supported by National Science Foundation grants 2138259, 2138286, 2138307, 2137603, and 2138296.

## Supplemental Information

**Figure S1:**
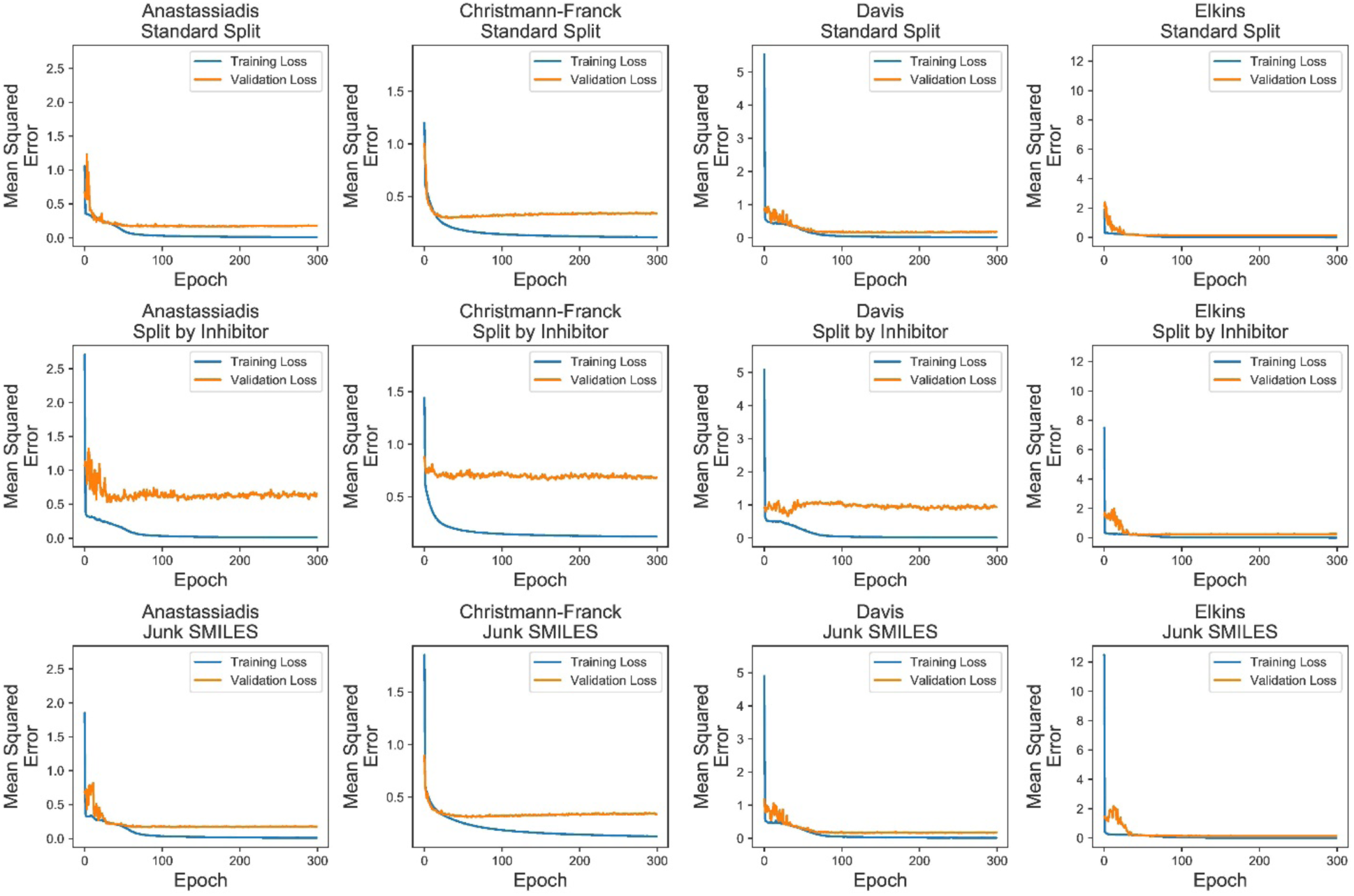
Training Set and Validation Set Loss Plots for Neural Network Model. We use the settings with lowest validation loss in the further evaluation of our model.

**Figure S2:**
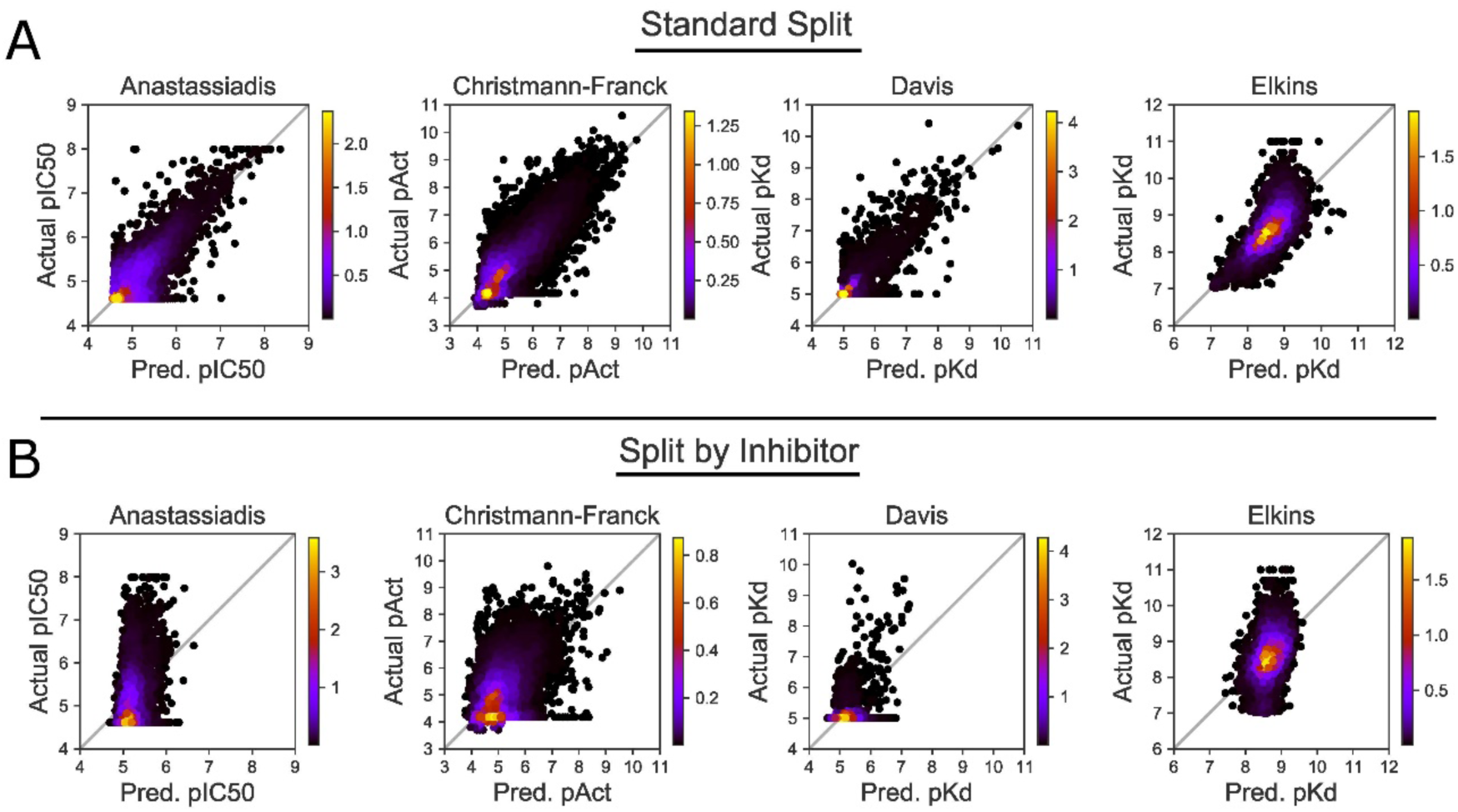
Using “Split by Inhibitor” diminishes model performance relative to “Standard Split”. These scatterplots correspond to the same data shown in Figure 3 as density plots.

**Figure S3:**
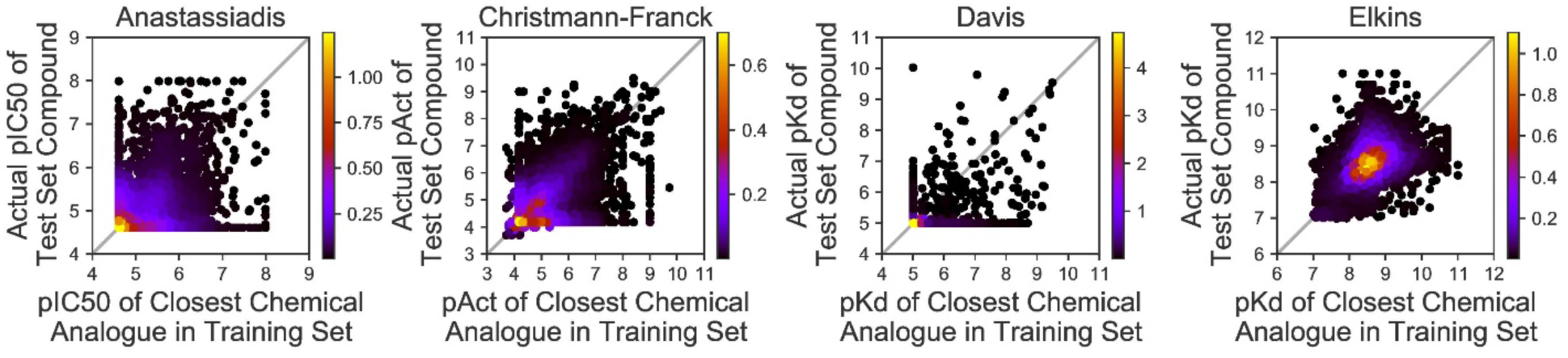
Using a tokenized inhibitor encoding does not diminish model performance. These scatterplots correspond to the same data shown in Figure 4 as density plots.

